# MiniScrub: *de novo* long read scrubbing using approximate alignment and deep learning

**DOI:** 10.1101/433573

**Authors:** Nathan LaPierre, Rob Egan, Wei Wang, Zhong Wang

## Abstract

Long read sequencing technologies such as Oxford Nanopore can greatly decrease the complexity of *de novo* genome assembly and large structural variation identification. Currently Nanopore reads have high error rates, and the errors often cluster into low-quality segments within the reads. Many methods for resolving these errors require access to reference genomes, high-fidelity short reads, or reference genomes, which are often not available. De novo error correction modules are available, often as part of assembly tools, but large-scale errors still remain in resulting assemblies, motivating further innovation in this area. We developed a novel Convolutional Neural Network (CNN) based method, called MiniScrub, for *de novo* identification and subsequent “scrubbing” (removal) of low-quality Nanopore read segments. MiniScrub first generates read-to-read alignments by MiniMap, then encodes the alignments into images, and finally builds CNN models to predict low-quality segments that could be scrubbed based on a customized quality cutoff. Applying MiniScrub to real world control datasets under several different parameters, we show that it robustly improves read quality. Compared to raw reads, *de novo* genome assembly with scrubbed reads produces many fewer mis-assemblies and large indel errors. We propose MiniScrub as a tool for preprocessing Nanopore reads for downstream analyses. MiniScrub is open-source software and is available at https://bitbucket.org/berkeleylab/jgi-miniscrub

## 1 Introduction

Long read sequencing has become increasingly important in recent years, with sequencing technologies from companies such as Pacific Biosciences [1] and Oxford Nanopore [2] seeing wide use in a variety of applications including genome assembly [1, 10], detection of antimicrobial resistance genes [18], sequencing personal transcriptomes [19], and improving draft genomes [20]. Sequence assembly is one of the most promising and widely-explored of these applications, as long repeat sections have been shown to be among the most important factors that affect assembly quality [12, 13], and long sequencing reads are much more capable of resolving these long repeats. Theoretical analysis has indicated that increasing read length from 100bp to 1000bp significantly simplifies the *de Bruijn* graphs used in assembly algorithms and can increase N50 size by six folds [12].

However, current single molecule, long sequencing reads also have very high error rates, ranging from 5% to 40% [10] per read and often average about 10% to 20% [1, 9] up to as high as 30–40% [10, 21], depending on variables such as the type and version of the sequencing technology and the experiment being performed. These high error rates makes assembly and other applications inefficient or error-prone [10, 9, 21, 2] and it is thus critical that methods be developed towards addressing this issue so that the potential of long read sequencing can be fully realized. Many current solutions involve “hybrid error correction” [9–11] by performing an additional sequencing run using low-error short reads and aligning them to the long reads, followed by a consensus approach to produce the correct sequence. Despite their success [9–11], the requirement for extra sequencing runs, often with different technologies, imposes additional monetary and temporal burdens [22]. Another approach involves re-analyzing the raw signal output by the sequencing machines to call the correct bases in the reads [23, 24], but researchers may not have this raw signal data available, and these methods are generally optimized for a specific version of a certain sequencing technology, and are thus not generally applicable and can become outdated quickly [25].

Thus, it is desirable to have a completely *de novo* method for improving long sequencing reads that does not rely on any information other than the reads themselves and is generally applicable across many technologies. Gene Myers [7] and others [10] observed that long read errors tend to locally cluster into certain low-quality “junk” segments, raising the possibility of “scrubbing” [7] (removing) these low-quality segments to significantly improve read quality. Recent work has addressed a related problem of *de novo* read error correction [28, 29]. However, even the best methods still produce quite a few misassemblies, suggesting independent methods are necessary for further improving assembly results.

Here we describe MiniScrub for *de novo* long Nanopore read scrubbing. MiniScrub performs read-to-read mapping to obtain alignments, converts them into images, followed by machine learning to identify the low-quality read segments to be scrubbed. We overcame several challenges inherent in this process. First, read-to-read alignment is a quadratic problem that traditional alignment tools such as BWA and Bowtie are not built to handle efficiently [8]. Second, because the dominant type of error in some long read sequencers is (potentially large) indels [2], exact alignments can be difficult to achieve. A recent method called MiniMap [4] addresses both of these problems by performing approximate read-to-read alignment by identifying read pairs that share a number of co-linear k-mers called “minimizers” [3, 4]. This avoids the difficult problem of exact mapping and runs over 50 times faster than BWA, making read-to-read mapping tractable [4]. Finally, because these read alignments are approximate, meaning that we only know the subset of k-mers that matched between reads, we are faced with a challenging pattern recognition problem. Namely, how many k-mers in a region of a given query read need to be supported by other reads, and by *how many* other reads, for that region of the query read to be considered high-quality?

We addressed this challenge by using deep learning, a recent machine learning paradigm [26]. Deep learning has been increasingly applied in recent years to problems within the biological sciences. A recent notable example is DeepVariant, which achieved superior results in variant calling competitions and benchmarks using a deep learning method called Convolutional Neural Networks [5]. In MiniScrub, we developed a novel method for encoding read-to-read mappings into “pileup” images, with information such as minimizers matched, quality scores, and distance between minimizers encoded in the color pixels of the images. These images were used as input into a Convolutional Neural Network (CNN), which is optimized to detect local patterns such as those present in images [26], to predict which read segments are of low-quality. See the Methods section below for an explanation of these terms. We show in the Results section that scrubbing with MiniScrub is able to robustly improve read quality and downstream assembly quality, even though assemblers already implement a read error correction step.

## 2 Materials and Methods

### 2.1 Method Overview

The three steps involved in MiniScrub are illustrated in Figure 1 and explained in further detail in the subsections below.

**Fig. 1:**
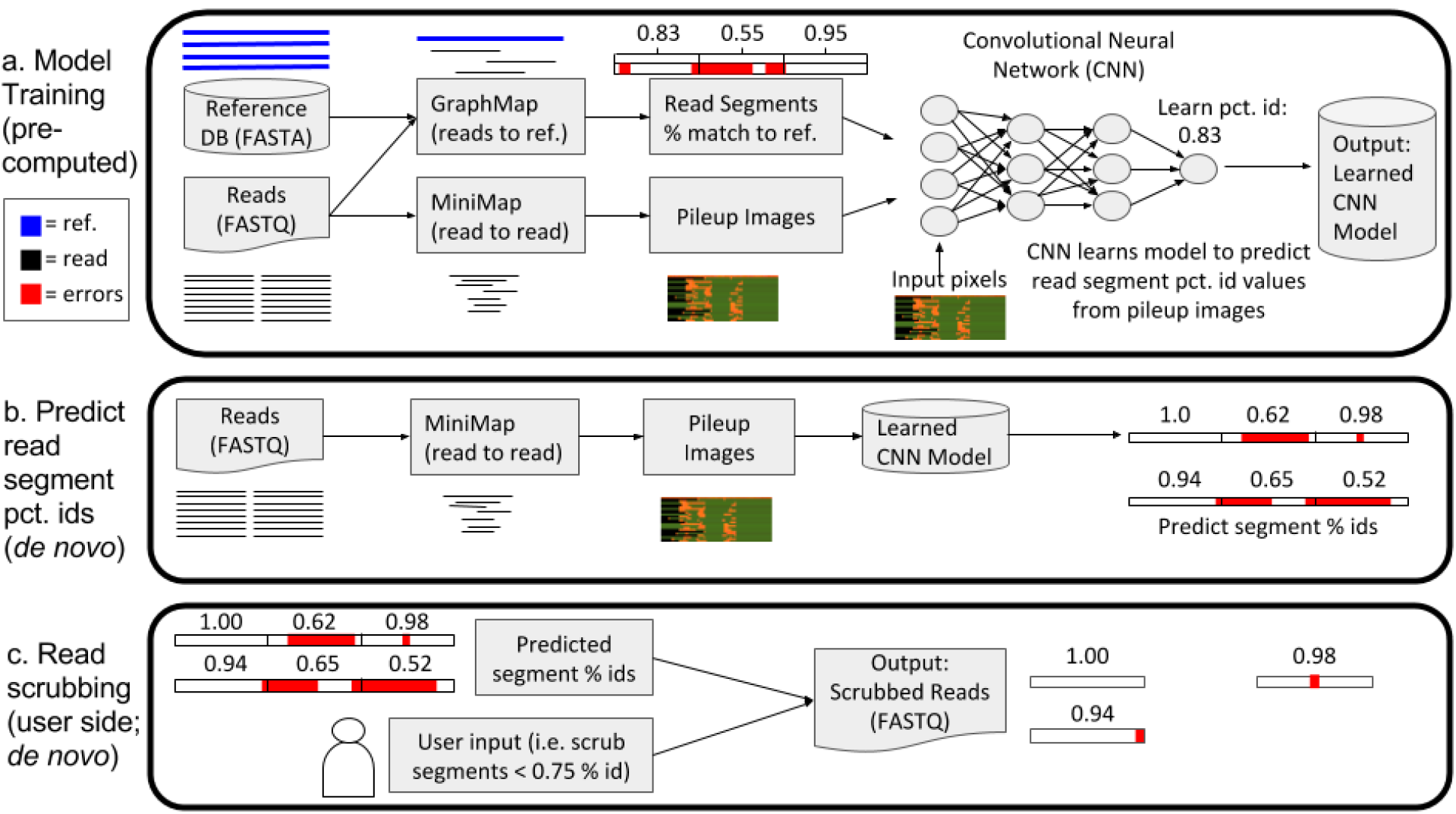
Overview of MiniScrub. As a precomputation step, the Convolutional Neural Network (CNN) must be trained to predict sequence segment percent identity (percent match to reference) from the read-to-read mappings. To generate ground-truth percent identity for read segments, reads are generated from known genomes in a reference database, then GraphMap [27] is used to map those reads to the reference, from which we calculate the percentage of bases from each read segment that match the reference genome. We also use MiniMap to generate *de novo* read-to-read mapping, then encode the information into an RGB “pileup” image for each read, which is then split up into shorter segments for better scrubbing resolution. We then train the CNN to learn the segment percent identity from the pileup images and save the model. On the user side, users run MiniMap on their set of reads and specify a cutoff threshold for read segments to scrub. The learned CNN model then predicts read segment percent identity and scrubs the segments below the quality threshold, outputting a new FASTQ file with the scrubbed reads.

The first step is training a CNN model, a “precomputation” step because it only needs to be done once for each setting of parameters and sequencing technology. The learned model can then be applied to any dataset, *de novo*, of the same sequencing technology that it was trained on. The model training step starts with building a training set with reads from a known reference genome. These reads are mapped using GraphMap [27]) to the reference genomes. We then divide a mapped read into short segments, defined by a number of minimizers (see following section). For each read segment we calculate its percent identity, e.g. the percentage of bases in the read that match the reference, as labels. We then use a modified version of MiniMap [4] to obtain read-to-read alignments between all reads in the training set (see below for details), and embed relevant alignment information (minimizers matched, distance between minimizers, and quality scores) into Red-Green-Blue (RGB) pixels to form “pileup” images. One image is generated for each read, and is then broken into the same short segments as above. A CNN model is then trained with the above data, learning a mapping from a pileup image of a read segment to the percent identity of that read segment. This process is explained more in the subsections below.

After a CNN model has been trained, users can use MiniScrub to generate images and segements of reads from the same sequencing technology, and predict the percent identity of each read segment. Finally, users can scrub out the segments below a user-set percent identity threshold (e.g. 0.8). Taking a FASTQ file as input, reads are split after low quality segments are removed, and they are written into a new FASTQ file.

### 2.2 Approximate Alignment Using MiniMap and Minimizers

Read-to-read mapping is generally not practical due to the quadratic nature of the problem (aligning all pairs of reads). Additionally, for some long read sequencing technologies such as Oxford Nanopore, (potentially large) indels are the dominant form of errors [2], making exact read-to-read mapping difficult. Here we use MiniMap to rapidly obtain approximate alignments between all read pairs [4] as it is efficient and robust to indels. MiniMap is based on identifying read pairs that share many co-linear “minimizers” [3]. Briefly, minimizers are the k-mers out of a set of *w* consecutive *k*-mers that minimize a certain function (such as alphabetical order). Minimizers have the benefit of guaranteeing that if two reads share the same *w* consecutive *k*-mers, they are guaranteed to share the same minimizer at that position; thus the minimizers shared between reads are an effective compressed representation of how closely reads match each other. We modified the MiniMap program to output the positions of all minimizers of all pairs of reads. Intuitively, if a minimizer in a given read is supported by many other reads, then there is a high likelihood that those k bases covered by the minimizer are error-free, while if no other reads covering the same sequence share that minimizer, it is likely to contain an error. For more details on minimizers, see the original paper by Roberts et. al [3].

### 2.3 Pileup Image Generation and Deep Learning with CNNs

Since CNNs are best adapted for image input, we developed a method for generating images from the read alignments, which we refer to as “pileup” images (following Gene Myers [7]). One “pileup” image was generated for each sequencing read, since MiniMap uses each read as a “reference read” once and gathers a set of “matching reads” for each reference read (forming a read “pile”). We randomly choose 24 of the matching reads (including the reference read itself) to generate the pileup image; we observed little gain in performance with more reads.

An example pileup image can be seen in Figure 1. Pileup images are generated by embedding the alignments between a reference read and its matching reads into Red-Green-Blue (RGB) pixels, forming an image. In the image, each column of pixels represents a minimizer in the reference read. The top row in each image represents the reference read, while subsequent pixel rows represent matching reads, thus each image has 24 rows. For each pixel, the red channel indicates whether or not a read contains this minimizer (yes: value 255, no: value 70). The green channel is the average base quality score doubled such that it ranges from 66–254. The blue channel represents the distance to the next minimizer; intuitively, if the blue pixel value is highly different between the reference read and a matching read, one of them likely has an indel. Finally, a (0,0,0) (black) pixel was entered for a section of a matching read that MiniMap did not identify as being part of the match. After the pileup image is generated for a read, it is divided into 48-minimizer-wide segments (segments of the reference read spanning 48 minimizers), meaning each image is 48 pixels long. This value was chosen for a strong balance between resolution and accuracy of predictions, but can be modified by the user.

For training CNN models, we use a modified version of VGG16, named after the Visual Geometry Group at Oxford and the number of layers in the network [6]. We chose VGG16 because it is among the most successful CNN architectures available [30], its architecture is open source [6] and widely implemented, and we view it as general-purpose and not overly-adapted to its original image classification task. The original architecture consists of 13 convolutional layers and three fully-connected layers. Each convolutional layer uses 3×3 pixel filters. VGG16 was originally developed to classify an image as belonging to one of 1000 categories, but since we are seeking to predict a real number from 0 to 1 (percent identity), we modified the VGG16 architecture to output a single real value. While we adapted the VGG16 architecture, we trained our own model weights from scratch, as we found the open-source VGG weights to be too adapted to their original image classification task to work well for our purposes.

### 2.4 Datasets and computing environments

We evaluated the performance of MiniScrub on two Oxford Nanopore datasets, which we refer to as the “Low Complexity” or “LC” dataset and the “High Complexity” or “HC” dataset. The LC dataset is used in most of our analyses, while the HC dataset is used in this section to evaluate cross-dataset performance. The LC dataset consists of two species sampled at high coverage, *E. coli* (204x coverage) and *S. koreensis* (140x coverage). In total, the LC dataset contained 747,598 reads averaging 2.6kb in length, out of which 724,140 were successfully mapped to the reference genomes; the rest were regarded as low quality and excluded from further anaylsis. The HC dataset consists of 260,930 reads sampled from 26 different species, at a much lower coverage (0.005x to 64x). The composition of the HC dataset is explained further in [34] and both datasets are available via the BitBucket repository linked in the abstract. Both datasets were sequenced with Oxford Nanopore MinION flowcell FLO-MIN107 and were basecalled with Albacore version 1.2.1.

The hardware used in the study was an NVIDIA DGX-1 deep learning system, which has 8 Tesla V100 GPUs, 128GB GPU Memory, 512GB System memory, 40,960 CUDA cores, 5,120 NVIDIA Tensor Cores, and a Dual 20-Core Intel Xeon E5–2698 v4 2.2 GHz Processor. However, only a small fraction of these resources were ultimately needed by our experiments, and GPUs are not required to run MiniScrub, though MiniScrub will be much slower without them. All experiments were performed with MECAT version 1.3, Canu version 1.7, TensorFlow version 1.8, and Keras version 2.2, with the exception of one Canu run with version 1.6, noted in the results section.

## 3 Results

### 3.1 MiniScrub robustly predicts low-quality segments within Nanopore reads

MiniScrub predicts the “percent identity” (percent of correct bases) of each segment of a read (defined by a number of bases or minimizers) and scrubs out segments below a user-set threshold, splitting the reads at the low-quality regions. To evaluate its performance, we use the Mean Squared Error and Pearson and Spearman correlations between the predicted percent identity by MiniScrub and the actual percent identity recovered from mapping the reads to the reference. Given our suggested user cutoff of 80% identity (or 0.8), we also calculated the sensitivity and specificity of MiniScrub’s ability to retain high-quality segments. In this case, high sensitivity translates into a low false negative rate, which is desirable as we should retain the high-quality segments as much as possible.

First, we evaluated MiniScrub’s performance by training its model on 25,000 reads, for 5 epochs, from the LC dataset (Methods) and tested its performance on the remaining reads. The results indicated that MiniScrub accurately predicted percent identity of read segments, with a Mean Squared Error of 0.003 and Pearson/Spearman correlation of 0.827/0.805 between the predicted and actual percent identities. Furthermore, given a user-specified cutoff of 0.8, MiniScrub had 95% sensitivity and 68.1% specificity, meaning that it retained 95% of read segments that were actually above the 0.8 threshold and successfully removed 68.1% of those below. This is a conservative setting, and more cutoff parameters can be tuned to scrub more aggressively (see Appendix for an exploration of this parameter).

We next assessed the performance of MiniScrub using two datasets generated from the same sequencing technology using the above metrics, to ensure that MiniScrub does not overfit to a single dataset. In contrast to the highly-covered, low-complexity community of *E. Coli* and *S. Koreensis* in the LC dataset, the HC mock community consists of 26 species at much lower average coverage, representing a very different application setting (Methods). We tested four different settings: training MiniScrub on the LC data and testing on the LC data, training on LC and testing on HC, training on HC and testing on LC, and training on HC and testing on HC. We ran MiniScrub for each setting by training the CNN on 25,000 images from the training dataset for 5 epochs, and calculated the mean squared error, Pearson correlation, Spearman rank correlation, and sensitivity/specificity at a 0.8 cutoff threshold on 5,000 images randomly drawn from the testing dataset. These results are shown in Table 1; note that the first column corresponds to the experiment described in the previous paragraph.

**Table 1:** Results from training and testing on different datasets. “LC” is a low complexity, high coverage (140x to 204x) community derived from 747,598 reads from only two species, *E. coli* and *S. koreensis*. “HC” is a high complexity, low coverage (0.005x to 64x) community derived from 260,930 reads from 26 species, described in [34]. The cutoff point for the sensitivity/specificity results was set at 0.8. We use the notation “LC train → HC test” to mean training the model on the LC data and testing it on the HC data.

|  | LC train → LC test | LC train → HC test | HC train → LC test | HC train → HC test |
| --- | --- | --- | --- | --- |
| Mean Sq. Error | 0.00300 | 0.00447 | 0.00312 | 0.00391 |
| Pearson | 0.827 | 0.747 | 0.809 | 0.772 |
| Spearman | 0.805 | 0.795 | 0.778 | 0.802 |
| Sensitivity | 0.950 | 0.891 | 0.938 | 0.889 |
| Specificity | 0.681 | 0.734 | 0.681 | 0.751 |

MiniScrub performs well not only within each dataset, but also across different datasets. Spearman correlation is similar across all settings, while models tested on the HC data trade off some sensitivity for higher specificity and have slightly worse Mean Squared Error and Pearson correlation. The small difference is likely due to the presence some low-coverage genomes in the HC data, as a small number of low-coverage reads will be less discriminatively scrubbed because they have less support from other reads. Despite this, the overall numbers are comparable, suggesting that MiniScrub recognizes the error patterns shared by very different datasets.

We also evaluated MiniScrub under various parameter settings, such as the number of matching reads compared with each query read and the length of the minimizer k-mers to use, and found that MiniScrub is robust to reasonable changes in parameter settings. Results are shown in the Appendix.

### 3.2 Scrubbing enriches the high-quality read poplulation

To test whether or not scrubbing improves read quality, we compared the reads from the LC dataset (Methods) before and after scrubbing by aligning them to the reference genome to obtain percent identity. As shown in Figure 2, after scrubbing we observed significant improvements in the read quality. First of all, the majority of the reads with a percent identity between 60–80 have been scrubbed, resulting in more, shorter reads between 85–95 percent identity. Even though MiniScrub does not perform error-correction, scrubbing out a small percentage of low-quality regions (presumably chimera junctions or indels) nevertheless raises average read percent identity by over 3% (from 83.1% to 86.2%). MiniScrub does not seem to perform much false scrubbing, as high quality raw reads (particularly those at 90% or higher accuracy) remain almost entirely intact.

**Fig. 2:**
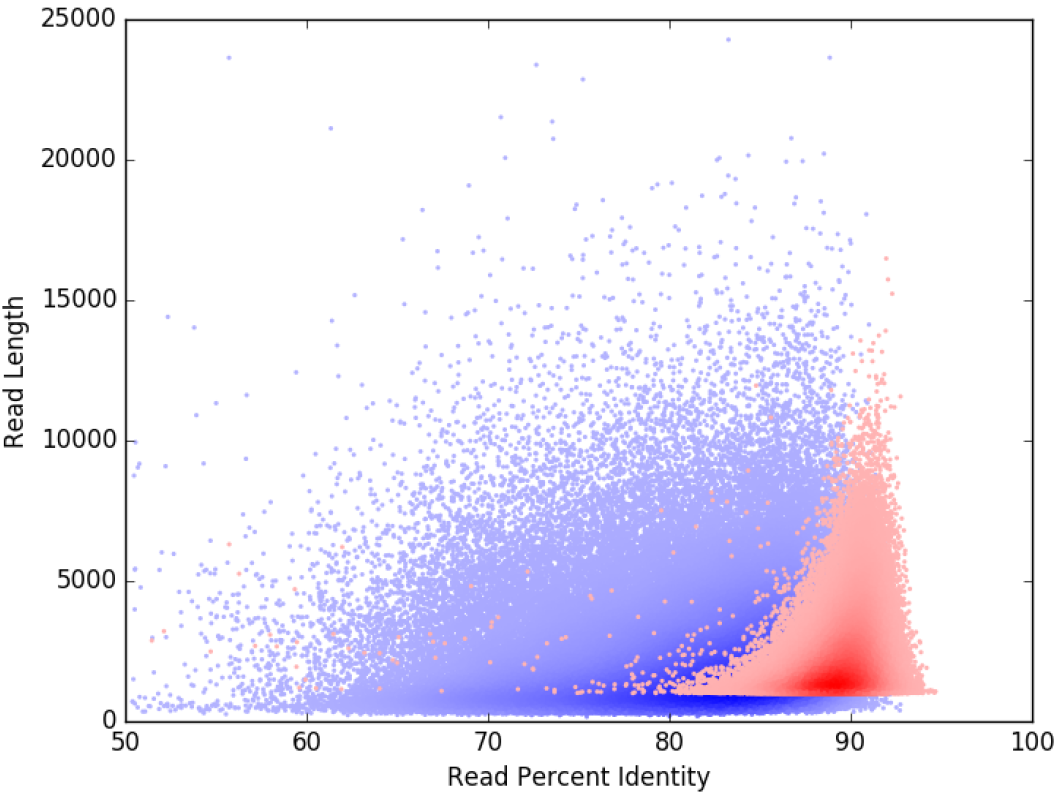
Density scatter plot showing average read quality improvement by MiniScrub versus raw reads. The X-axis shows read percent identity to the reference while the Y-axis shows read length. Raw reads are in blue while scrubbed reads are in red. The darkness of the color indicates increased “density” – more reads fall into a darker region of the graph than the lighter areas. MiniScrub scrubs out most of the low-quality segments in low quality reads while leaving high quality reads intact, increasing average read percent identity by over 3%, from 83.1% to 86.2%. Average read length decreased from 2.6kb to 1.1kb due to splitting reads where low-quality segments were removed.

**Fig. 3:**
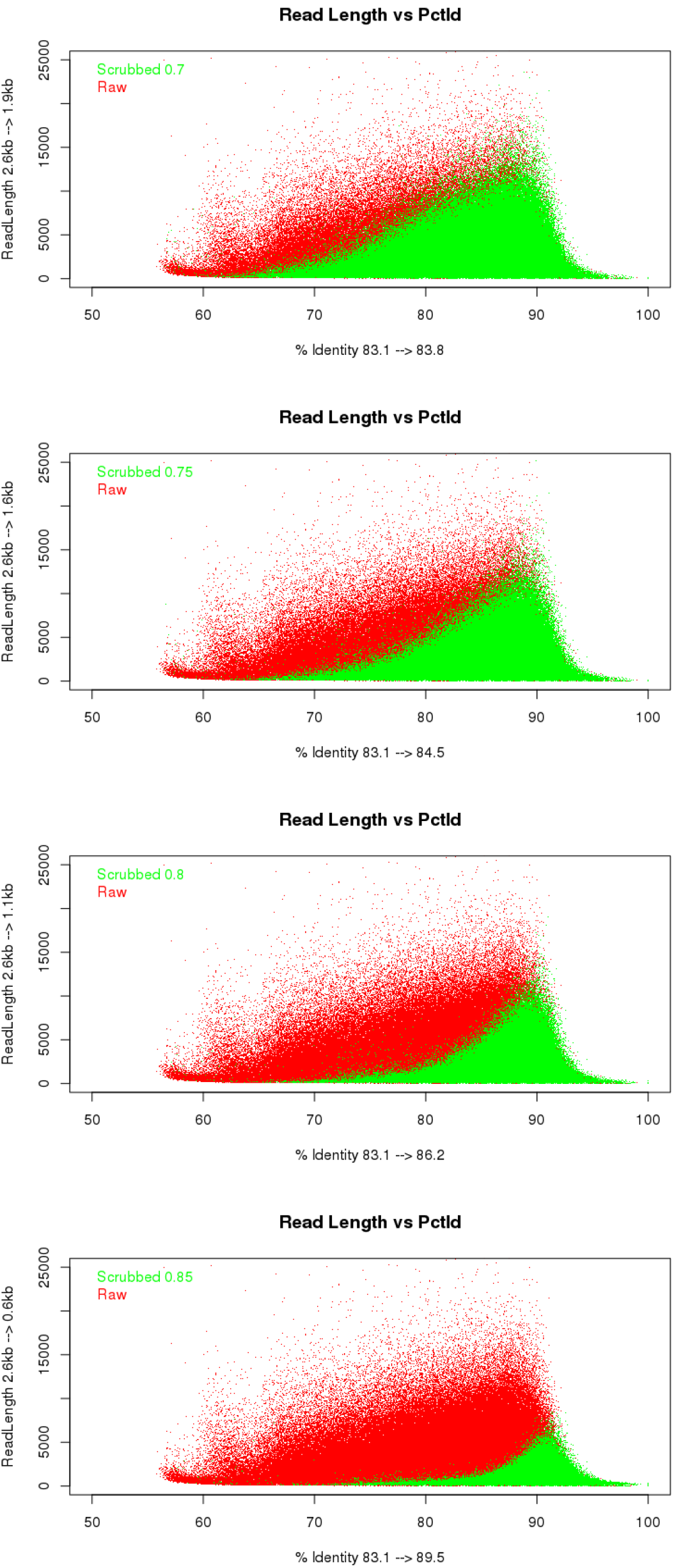
Plot of read lengths (x axis) versus read percent identity to the reference (y axis). Green is after scrubbing, red is before. From top to bottom, the cutoffs used are 0.7, 0.75, 0.8, 0.85.

### 3.3 MiniScrub leads to improvements in speed and/or accuracy of *de novo* assembly

We tested whether or not MiniScrub can be used as a preprocessing step to improve *de novo* assembly. Several recent long read assembly methods for Nanopore have been developed, including Canu [28], MECAT [31], DALIGNER [8], and more. We chose Canu (version 1.6) and MECAT (version 1.3) for this experiment, as Canu is a popular and well-established method, while MECAT is a newer method that is similar to Canu except with one of the slowest steps of Canu optimized to be faster [31].

We assembled the LC dataset with MECAT and Canu using either raw reads or scrubbed reads. MECAT seems to have problems with very long reads, so we split raw reads longer than 100kb into 100kb segments for it to run without errors. Results were evaluated using Quast [32] and are shown in Table 2.

**Table 2:**
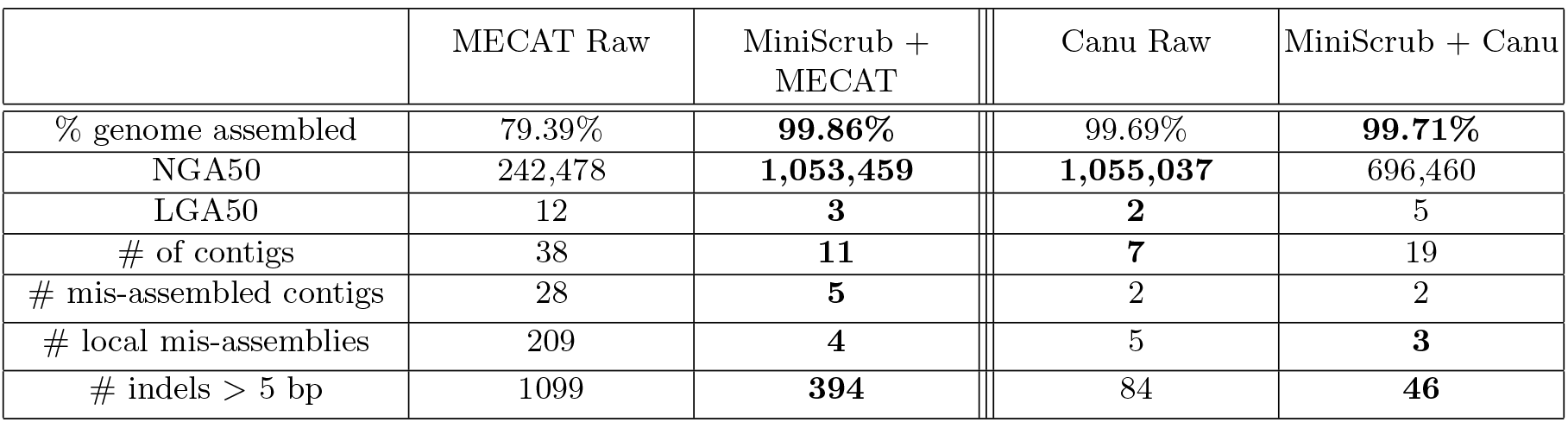
MiniScrub significantly improves assembly, tested with MECAT [31], increasing genome coverage and NGA50 while limiting LGA50, mis-assemblies, mismatches, and indels. Canu’s assembly had slightly reduced errors and misassemblies when reads were preprocessed with MiniScrub, but the assembly was more fractured, likely due in part to resolving large misassemblies and indels. Notably, Canu assembly of raw reads took about 3.5 days, while the MiniScrub+Canu pipeline took about 9 hours, likely due to a reduction in the amount of error correction needed in the latter situation. Results were evaluated using QUAST [32].

|  | MECAT Raw | MiniScrub +<br>MECAT | Canu Raw | MiniScrub + Canu |
| --- | --- | --- | --- | --- |
| % genome assembled | 79.39% | **99.86%** | 99.69% | **99.71%** |
| NGA50 | 242,478 | **1,053,459** | **1,055,037** | 696,460 |
| LGA50 | 12 | **3** | **2** | 5 |
| # of contigs | 38 | **11** | **7** | 19 |
| # mis-assembled contigs | 28 | **5** | 2 | 2 |
| # local mis-assemblies | 209 | **4** | 5 | **3** |
| # indels > 5 bp | 1099 | **394** | 84 | **46** |

After scrubbing the reads, MECAT assembly quality was dramatically improved, with genome coverage increasing from about 79.39% to 99.86%, mis-assembled contigs decreasing from 28 to 5, local mis-assemblies decreasing from 209 to 4, and the number of indels longer than 5bp reduced from 1099 to 394. NGA50 and LGA50 measure the size and number of correctly-assembled contigs required to cover half of the reference genome, with contigs taken in descending order by length. Concretely, MECAT assembly with raw reads required 12 contigs with size 242,478bp or longer to cover half of the reference genome, while assembly with the scrubbed reads only required 3 contigs, which were all 1,053,459bp or longer. Thus, the scrubbed reads produce an improved assembly with fewer, longer contigs that have fewer mis-assemblies and cover much more of the reference genome. Notably, MECAT applies an error correction step [31], so MiniScrub significantly improves performance as a preprocessing step even when subsequent read error correction is performed. This illustrates the potential of using read scrubbing, read error correction, and assembly in tandem.

The difference between Canu assemblies with raw reads and scrubbed reads is much smaller compared with MECAT assemblies. Scrubbing reduces local misassemblies from 5 to 3, and from 84 large indels to 46, while the assembly becomes slightly more fragmented.

However, Canu runtime was dramatically reduced on scrubbed reads, going from over 3.5 days on raw reads to 9 hours on scrubbed reads, including the read scrubbing step. In contrast, MECAT was much faster, taking about 2.5 hours with raw reads, but about 9 hours to scrub the reads and run assembly. This suggests that the dataset could be assembled quickly and accurately using either MiniScrub+MECAT or MiniScrub+Canu, but without scrubbing the assembly could be inaccurate or time-consuming.

### 3.4 MiniScrub versus other alternative preprocessing tools

There are two existing tools, Porechop^5^ and NanoFilt [33], that have similar capability to improve Nanopore read quality. We therefore compared their performance with MiniScrub as preprocessing methods for assembly.

It has been reported that 50x sequencing coverage of a single genome is a good setting for assembly [28]. To prepare this setting, we randomly selected reads from the LC dataset that mapped to the *E. coli* genome, up to 50x coverage. These reads were preprocessed by MiniScrub, Porechop, or NanoFilt before they were assembled by MECAT or Canu. Quast [32] metrics were used for assembly quality evaluation, and the results are shown in Table 3.

**Table 3:**
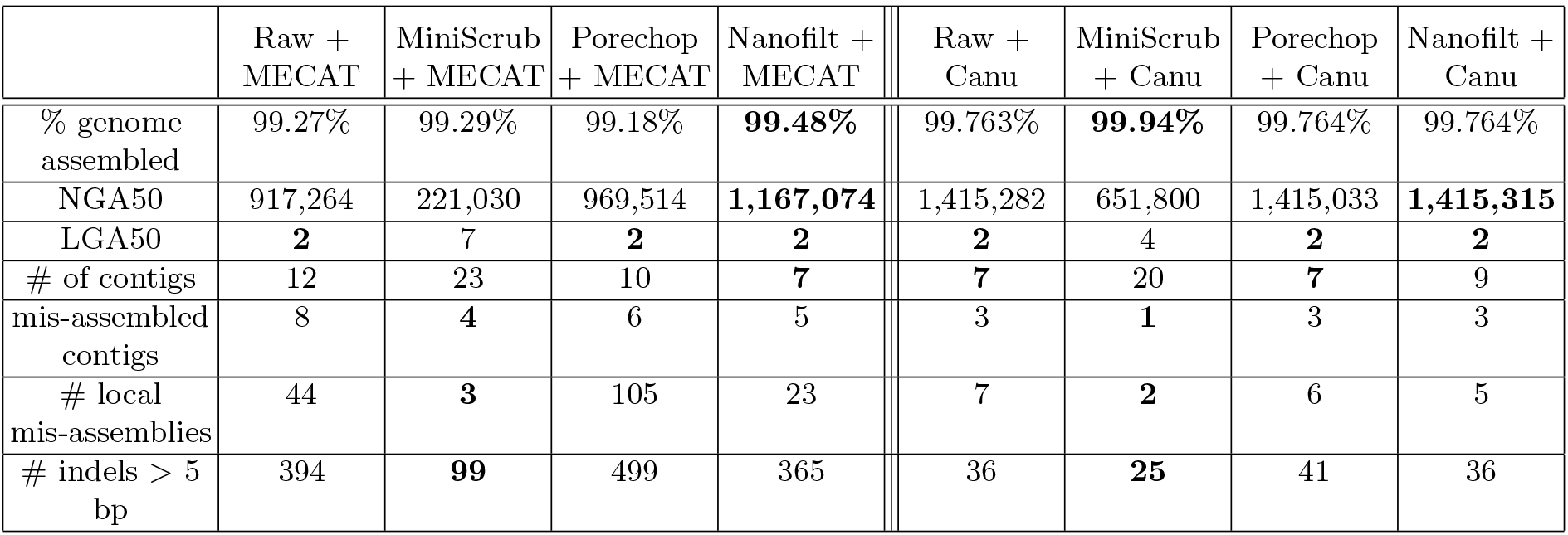
Comparing the performance of different Nanopore read preprocessing tools on a *E. coli* dataset with 50x coverage. Reads were preprocessed by MiniScrub, Porechop and nanofilt and then assembled using MECAT or CANU. Quast metrics were used for evaluation.

|  | Raw +<br>MECAT | MiniScrub<br>+ MECAT | Porechop<br>+ MECAT | Nanofilt +<br>MECAT |  | Raw +<br>Canu | MiniScrub<br>+ Canu | Porechop<br>+ Canu | Nanofilt +<br>Canu |
| --- | --- | --- | --- | --- | --- | --- | --- | --- | --- |
| % genome<br>assembled | 99.27% | 99.29% | 99.18% | **99.48%** |  | 99.763% | **99.94%** | 99.764% | 99.764% |
| NGA50 | 917,264 | 221,030 | 969,514 | **1,167,074** |  | 1,415,282 | 651,800 | 1,415,033 | **1,415,315** |
| LGA50 | **2** | 7 | **2** | **2** |  | **2** | 4 | **2** | **2** |
| # of contigs | 12 | 23 | 10 | **7** |  | **7** | 20 | **7** | 9 |
| mis-assembled<br>contigs | 8 | **4** | 6 | 5 |  | 3 | **1** | 3 | 3 |
| # local<br>mis-assemblies | 44 | **3** | 105 | 23 |  | 7 | **2** | 6 | 5 |
| # indels > 5<br>bp | 394 | **99** | 499 | 365 |  | 36 | **25** | 41 | 36 |

Consistent with the above results, MiniScrub greatly reduces the number of misassembled contigs, local misassemblies, and long indels while slightly decreasing assembly contiguity, likely due in part to breaking long reads that contain errors. In contrast, Porechop led to an increase in local misassemblies on the MECAT assembly and long indels on both datasets. Nanofilt helped reduce some mis-assemblies but not large indels. This extra reduction in assembly errors comes with a small time cost, as Nanofilt took 1 minute and 45 seconds to run, Porechop took 3 minutes and 39 seconds, while MiniScrub took 26 minutes and 14 seconds.

## 4 Discussion

We developed a method called MiniScrub that performs *de novo* long read scrubbing using the combined power of fast approximate read-to-read alignments, deep Convolutional Neural Networks, and a novel method for pileup image generation. We demonstrated that it accurately scrubs out low-quality segments within Nanopore raw reads to improve overall read quality, and the scrubbed reads lead to fewer assembly errors. In addition, MiniScrub outperforms alternative preprocessing tools in terms of leading to fewer misassemblies and large indels in *de novo* assembly.

Besides *de novo* genome assembly, we expect read scrubbing may also improve other downstream analyses, such as read error correction, large structural variation detection, etc. As MiniScrub uses a generic framework, it is possible that MiniScrub can learn technology-specific error profiles. Even though we focused on Oxford Nanopore reads in this study, read scrubbing may work on other long read technologies, such as PacBio SMRT. One would have to train a new CNN model for each different sequencing technology.

As MiniScrub splits reads at the point of scrubbing (chimera junctions or indels), spliting at indels will lead to lower assembly contiguity, especially affecting the low-coverage regions. Even though this may be a trade-off between contiguity and fewer errors, this leaves room for future improvements. One of the potential improvements would be to train the model to discriminate the chimera junctions and indels, and only split the chimeric reads while leaving those with large indels for read correction modules to fix.

In our current CNN model, both convolution and pooling are locally performed for small patches of the pileup images separately, without considering contextual dependencies between different patches. An interesting methodological direction would be to change our model to a Convolutional Recurrent Neural Network (CRNN) by adding Recurrent Neural Network (RNN) layers to learn contextual dependencies among sequential data through the recurrent (feedback) connections. This CRNN model may further enhance the predictive performance, especially the ability to detect low-quality regions.

## 5 Acknowledgements

The authors would like to thank NERSC, the NSF, and the NIH for their funding and support. The work conducted by Rob Egan and Zhong Wang was supported by the Office of Science of the U.S. Department of Energy under Contract No. DE-AC02- 05CH11231. The work conducted by Nathan LaPierre and Wei Wang was supported by NSF grant DGE-1829071 and NIH grants 5T32EB016640–05 and 4T32EB016640–04.

## Appendix A MiniScrub’s performance across different parameter settings

By default, MiniScrub has a default pileup image size of (Length, Depth) = (48, 24), meaning 48 minimizer-wide segments, and up to 23 matching reads for each query read. Additionally, MiniScrub uses minimizers with settings (w,k) = (5,15), meaning that a minimizer k-mer of length 15 is selected out of each 5 consecutive 15-mers. We sought to evaluate whether MiniScrub was effective under these default parameter settings, and whether it would be robust to reasonable adjustments to these parameters. Starting from the default settings of (Length, Depth) = (48, 24) and (w,k) = (5,15), we varied each pair of parameters in turn while holding the other pair constant. Namely, we evaluated the settings of (Length, Depth) = (36, 36) and (w,k) = (7,17). We ran MiniScrub for each setting by training the CNN on 25,000 images from the LC dataset for three epochs and calculating the mean squared error, Pearson correlation, Spearman rank correlation, and sensitivity/specificity at a 0.8 cutoff threshold. These results are shown in Table 4, along with results from the default parameters for comparison. As the table shows, MiniScrub performs robustly under all tested parameter settings, giving similar performance.

**Table 4:** Performance with different parameter settings. w and k refer to the minimizer parameters, while Length and Depth refer to the length and depth of each pileup image segment, which correspond to the number of minimizers in that read segment and the number of matching reads used. The default settings are (w,k) = (5,15) and (Length, Depth) = (48, 24). The columns show the performance when varying one of these settings and with cutoff 0.8.

|  | Default | (Length, Depth)= (36, 36) | (w,k)= (7,17) |
| --- | --- | --- | --- |
| Mean Sq. Error | 0.00300 | 0.00329 | 0.00305 |
| Pearson | 0.827 | 0.821 | 0.830 |
| Spearman | 0.805 | 0.786 | 0.810 |
| Sensitivity | 0.950 | 0.934 | 0.914 |
| Specificity | 0.681 | 0.693 | 0.780 |

## Appendix B Effects of Cutoff Threshold on Read Quality Improvement

Figure 3, on the following page, shows the read improvement effects of MiniScrub on the LC data (Methods) at various cutoff thresholds (0.7, 0.75, 0.8, 0.85) as compared with raw reads. As the cutoff threshold is increased, less overall data is retained, but the remaining data is higher-quality. This allows the user to customize MiniScrub to their needs. Concretely, scrubbing with a 0.7 cutoff retains upwards of 80% of the data, but only improves read percent identity by 0.7%, while scrubbing with a 0.85 cutoff retains less than half of the data but improves average read percent identity by about 6.5%.

## Footnotes

5 https://github.com/rrwick/Porechop

